# Single-cell and spatial transcriptome characterize coinhibitory cell-cell communications during histological progression of lung adenocarcinoma

**DOI:** 10.1101/2024.06.04.597379

**Authors:** Liu Hui, Qianman Gao, Judong Luo

## Abstract

Lung adenocarcinoma, a prevalent and lethal malignancy globally, is characterized by significant tumor heterogeneity and a complex tumor immune microenvironment during its histologic pattern progression. Understanding the intricate interplay between tumor and immune cells is of paramount importance as it could potentially pave the way for the development of effective therapeutic strategies for lung adenocarcinoma. In this study, we run comparative analysis of the single-cell transcriptomic data derived from tumor tissues exhibiting four distinct histologic patterns, lepidic, papillary, acinar and solid, in lung adenocarcinoma. Our analysis unveiled several co-inhibitory receptor-ligand interactions, including PD1-PDL1, PVR-TIGIT and TIGIT-NECTIN2, that potentially exert a pivotal role in recruiting immunosuppressive cells such as M2 macrophages and Tregs into LUAD tumor, thereby establishing immunosuppressive microenvironment and inducing T cells to exhaustion state. Furthermore, The expression level of these co-inhibitory factors, such as NECTIN2 and PVR, were strongly correlated with low immune infiltration, unfavorable patient clinical outcomes and limited efficacy of immunotherapy. Furthermore, we conducted immunofluorescence assay and spatial transcriptomic sequencing to validated the spatial co-localization of typical co-inhibitory factors. We believe this study provides valuable insights into the heterogeneity of molecular, cellular interactions leading to immunosuppressive microenvironment during the histological progression of lung adenocarcinoma. The findings could facilitate the development of novel immunotherapy for lung cancer.

## Introduction

Lung cancer has the highest incidence among all malignant tumors in men worldwide, causing over a million deaths every year. Lung adenocarcinoma (LUAD) is the predominant histological subtype [1], accounting for 40% of lung cancer patients. The progression and prognosis of LUAD are closely associated with the emergence of morphologically distinct tumor regions, termed histologic patterns [2], including lepidic, papillary, acinar, and solid patterns. These patterns reflect the increasing aggressiveness of the tumor as it progresses. Previous studies have reported that genomic alterations, such as tumor mutational burden (TMB) and copy number variation (CNV), also increased during the histological progression [3, 4]. However, the tumor heterogeneity and intercellular interactions that contribute to the immunosuppressive tumor microenvironment (TME) during the histological progression remain elusive. Solid tumors are often characterized by highly complex TME that comprises infiltrating immune cells, stromal cells, chemokines, and extracellular matrix components. Tumor cells dynamically interact with TME to create a low-oxygen, low-pH, pro-inflammatory, and immunosuppressive environment that promotes the progression of cancer and influences patient’s response to drug treatment. Single-cell RNA sequencing (scRNA-seq) enables a comprehensive analysis of the cellular diversity and heterogeneity within tumor tissues [5–7]. By profiling the gene expression of individual cells, scRNA-seq offers a valuable chance to reveal the intratumoral heterogeneity and cell-cell communication mediated by ligand-receptor interactions [8]. For instance, Qi et al. studied the intratumoral heterogeneity in colorectal cancer (CRC) tumors and adjacent tissues, with a particular focus on the interplay between SPP1+ macrophages and FAP+ fibroblasts. The co-localization of these two cell types was substantiated through the use of immunofluorescence staining and spatial transcriptomics [9]. Zhang et al. have studied the interaction between malignant cells and regulatory T cells (Tregs) in intrahepatic cholangiocarcinoma (ICC), and found that TIGIT-PVR signal was enriched between Tregs and malignant cells [10]. Kim et al. have studied the cell-cell communications in different metastatic tumors of lung adenocarcinoma [11]. Nevertheless, there was a dearth of studies exploring the molecular features and intercellular interactions during the histologic pattern progression of lung adenocarcinoma.

In this study, we employed the scRNA-seq data to elucidate molecular heterogeneity, intercellular communications, and immunosuppressive landscapes of lung adenocarcinoma with different histologic patterns. We revealed that the co-inhibitory ligand-receptor interactions were more prevalent and prominent in the solid pattern. Specifically, the tumors with high expression levels of NECTIN2 and PVR exhibited low immune infiltration level, limited response to immunotherapy and poor prognosis. Moreover, our immunofluorescence experiment confirmed the spatial co-localization of cells expressing high levels of PVR and TIGIT molecules in the solid pattern, but not in the lepidic pattern. We further performed spatial transcriptome sequencing to validate the co-localization of these co-inhibitory factors. These findings suggested that tumor cells escaped immune surveillance by upregulating the expression of NECTIN2 and PVR immune inhibitory molecules to reduce immune cell infiltration. As far as our knowledge, this is the first study that used the single-cell and spatial transcriptomic data to explore the molecular mechanism underlying the histologic pattern progression of LUAD. We believe that our study provided insight into the immunosuppressive microenvironment formation and the underlying mechanism during the lepidic-to-solid progression of lung adenocarcinoma.

## Materials and Methods

### Ethics approval statement

The ethical approval has been obtained from the institutional review board the Affiliated Tongji Hospital of Tongji University, Shanghai, China. Patients gave informed consent at hospitalization.

### Single-cell transcriptomic data source

The scRNA-seq data of four histologic patterns was downloaded from a prior study [12], which includes one lepidic sample, one papillary sample, two acinar samples and two solid samples.

### Spatial transcriptomics library and sequencing

Two specimens of lung adenocarcinoma, each corresponding to the lepidic and solid histologic patterns as confirmed through histologic scrutiny, were procured in accordance with standard surgical protocols. These specimens underwent a process of formalin fixation and were subsequently encapsulated within paraffin-embedded tissue blocks. The specimens were then sectioned and subjected to hematoxylin and eosin (H&E) staining to facilitate subsequent imaging at a resolution of 40x (equivalent to 0.25 micron/pixel) via the use of Aperio GT450 scanners. The tissue slides were then conveyed to the Genomics core, where following the decoverslipping of the tissue, the Visium CytAssist device was employed to transfer transcriptomic probes from the original glass slides to capture areas on Visium slides measuring 11mm x 11mm. Comprehensive transcriptomic profiling was achieved post mRNA permeabilization, through poly(A) capture and probe hybridization. The resultant libraries were sequenced utilizing the Illumina Novaseq 6000, using paired-end sequencing with a read length of 150 base pairs.

### Spatial transcriptomics data analysis

Visium spatial transcriptomics profiles for samples contained 18,085 genes across several thousand locations throughout each slide. The profiles were then subjected to filtering process: spots with fewer than 500 genes, genes expressed in fewer than 3 spots, and spots with mitochondrial gene expression exceeding 15% were excluded from the analysis. Following the removal of regions lacking tissue using a custom annotation tool augmented by the SAM, a total of 4992 spots were obtained from the lepidic sample and 3701 spots from the solid sample. Each Visium spot covers a circular capture area with a diameter of 50-micron ( 200 pixels) at 40x magnification. Histology images and FASTQ files were processed using the Space Ranger pipeline and sequencing data were aligned to the human reference genome (Ensemble Genome GRCh38). The output generated was imported into Seurat for further analysis, including dimensional reduction, clustering and data integration.

### scRNA-seq data quality control and cell type annotation

The scRNA-seq data was processed and analysed using the Seurat R package. Cells were removed if they had more than 20,000 UMIs, more than 5,000 or fewer than 500 expressed genes, or >30% UMIs that were derived from the mitochondrial genome. The batch effect across different samples was mitigated using Seurat’s CCA algorithm. Prior to merging the data, the expression matrix of each sample was standardized using the NormalizeData function and the top 2000 variable genes were selected for principal component analysis (PCA) using the *FindVariableFeatures* function. Next, the merged data was normalized using the ScaleData function, clustering analysis was performed using the first 30 principal components at resolution 0.8, and visualization was performed using the UMAP algorithm. The classic marker genes used to identify the 13 main cell types were jointly determined by the FindAllmarkers function and literature research, and finally the following genes were used to identify different cell types: CD8^+^ T (CD8A, CD8B), CD4^+^ T (IL7R), Tumor cells (EPCAM, KRT19, KRT7), Tregs (FOXP3, IL2RA), Macrophages (CD68, C1QA, C1QC), Plasma cells (IGHG1, JCHAIN, CD79A), B cells (MS4A1, CD19), NK cells (NKG7, GNLY, KLRD1 ), Monocytes (S100A8, S100A9, FCN1), Fibroblasts (LUM, PDGFRA, ACTA2), Mast cells (TPSAB1, TPSB2, KIT, GATA2), Endothelial cells (VWF, CLDN5, CALCRL), Plasmacytoid dendrites Stem cells (LILRA4, IL3RA, CLEC4C).

### scRNA-seq-based CNV estimation

The inferCNV R package was used to infer copy number alterations in various cell types, using the gene expression of B cells as a reference. For parameter setting of inferCNV, cutoff was set to 0.1, cluster by groups was set to T. The cell hierarchical clustering method of the ward deviation sum of squares method (ward.D2) was used to cluster cells and denoised the results.

### Intercellular interaction analysis

CellPhoneDB was used to infer interactions between different cell types. For genes expressed in a cell population, the percentage of cells expressing the gene and the average gene expression were calculated, and the gene was removed if it was expressed in only 10% or less of the cells in the population. Cell-cell interactions were inferred from the gene expression levels by 1000 permutation tests, and then an adjacency matrix was generated for all cell-cell interactions and displayed on a heat map. The relative expression levels (z-scores) of partial ligands or receptors were visualized using the ggplot2[13] package.

### Immunofluorescence staining

The immunofluorescence staining was performed to examine the localization of TIGIT, NECTIN2 and PVR in the pathological tissues of patients with two histologic patterns. Paraffin sections of pathological tissues from LUAD patients with both histologic patterns were first deparaffinized and hydrated, followed by antigen retrieval. The primary antibody mixture was added at a ratio of 1:1000: TIGIT (Abcam, Ab243903) + NECTIN2 (Abcam, Ab269721) or TIGIT (Abcam, Ab243903) + PVR (Ab307687), placed in a wet box, and incubated overnight at 4°C; Then added two fluorescent secondary antibodies: Fluorescein (FITC)-conjugated Affinipure Goat Anti-Rabbit IgG(H+L), Proteintech, SA00003-2 + CoraLite594-conjugated Goat Anti-Mouse IgG(H+L), Proteintech, SA00013-3 and Fluorescein (FITC)-conjugated Affinipure Goat Anti-Rabbit IgG(H+L), Proteintech, SA00003-2 + Rhodamine (TRITC)-conjugated Goat Anti-Rat IgG(H+L), Proteintech, SA00007-7, placed in a humid box at room temperature and incubated in the dark for 2 hours; then counterstaining with DAPI and incubated in the dark for 15 minutes, and finally sealed the slides with an anti-fluorescence quencher, and dried them at 37°C for storage.

### Trajectory analysis

Slingshot was used to reveal the developmental trajectory of CD8^+^ T cells, and the identified pathways were mapped to UMAP projections for visualization.

### Survival analysis

The survival package was used to perform proportional hazards hypothesis tests and perform fitted survival regressions. The survminer[14] package was used to plot the Kaplan-Meier survival curves.

### Tumor immune microenvironment analysis

The CIBERSORTx tool was used to calculate immune cell infiltration levels using the gene expression matrix, the limma package was used for differential analysis of immune cells, and the vioplot package was used for visualization. The score of the TME was calculated using the ESTIMATE package, and the t-test was used to evaluate statistical significance. p<0.05 is represented by *, p<0.01 is represented by **, and p<0.001 is represented by ***.

### Correlation analysis

The Spearman correlation between overall survival and recurrence free survival on MSK-IMPACT LUAD cohort was calculated for lepidic and solid patterns, respectively. Spearman correlation coefficients were also calculated between biomarker genes on TCGA-LUAD cohort. The scatter plots were visualized using the ggplot2 R package.

## Results

### Different histologic patterns showed molecular heterogeneity and prognosis

Tumor heterogeneity arises from diverse molecular mechanism during tumor progression[15], resulting in substantial differences in the tumor microenvironment (TME), tumor mutation burden (TMB), and clinical outcomes. We explored the TMB differences between the four histologic patterns On the TCGA-LUAD (lepidic:n=10; papillary:n=47; acinar:n=65; solid:n=58) and MSK-IMPACT-LUAD (lepidic:n=88; papillary:n=43; acinar:n=368; solid:n=68) cohorts, and found that the solid pattern had higher TMB than the other patterns in both cohorts (Fig. 1a), indicating a higher rate of genetic mutations in the solid pattern. Moreover, On the MSK-IMPACT-LUAD cohort, we observed significant differences in the survival outcomes between four histologic patterns. Patients with solid pattern had lower recurrence-free survival (RFS) than those in other patterns (Fig. 1b), indicating higher degree of tumor growth and invasiveness in the solid pattern. We also found that OS was highly correlated with RFS in patients with lepidic pattern and only weakly correlated in those with solid pattern (Fig. 1c). This suggested that the tumor in solid pattern exhibited higher degree of resistance to drug treatment.

**Fig. 1:**
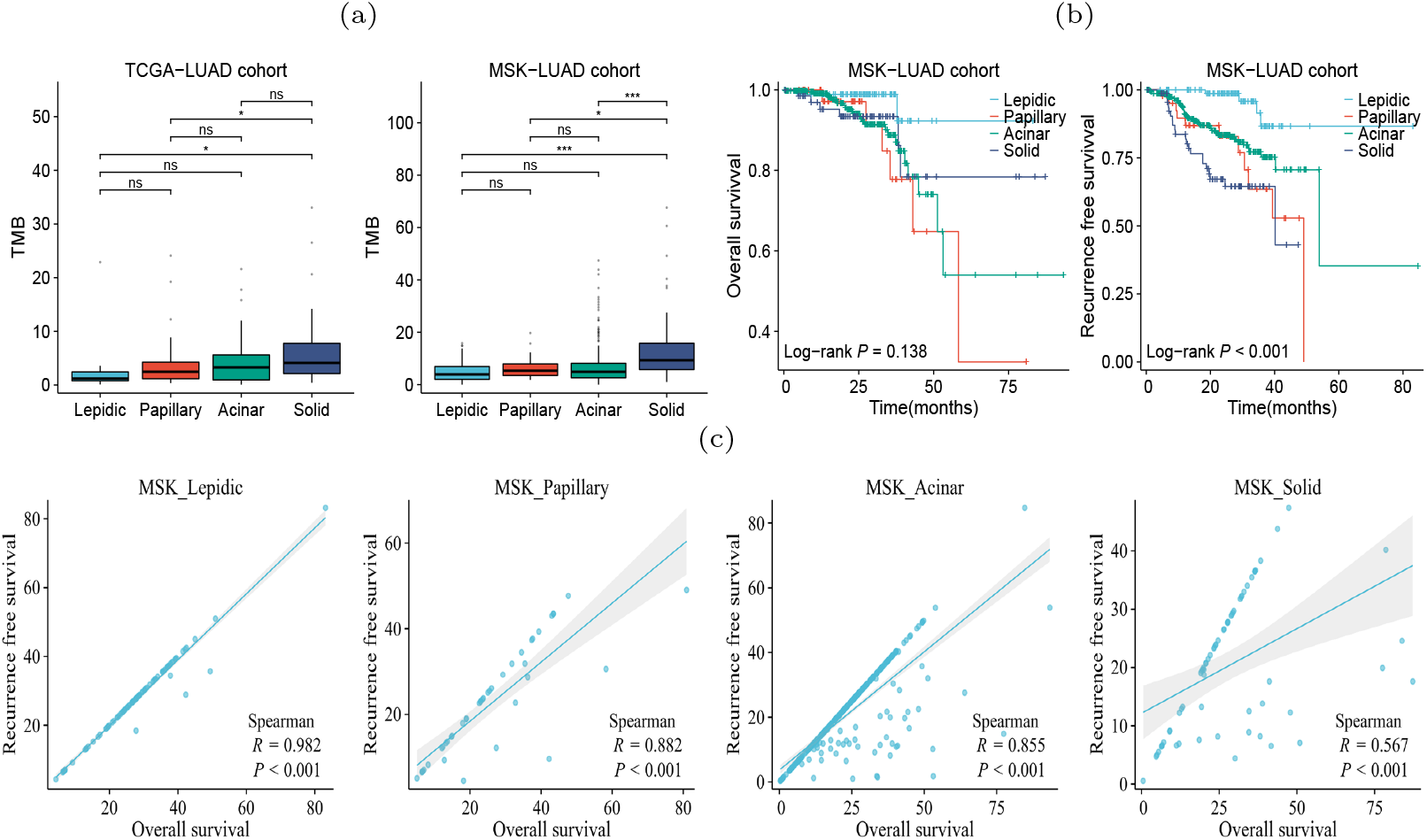
Heterogeneity of four histologic patterns of LUAD. (a) Differences in TMB between patient subgroups of four histologic patterns on TCGA and MSKCC cohorts. (b) Differences in OS and RFS between patient subgroups of four histologic patterns on LUAD in MSKCC cohorts. (c) Spearman correlation between OS and RFS of the patients with four histologic patterns on the MSKCC cohort. p<0.05 is represented by *, p<0.01 is represented by **, and p<0.001 is represented by ***.

### scRNA-seq revealed increasing tumor heterogeneity during histologic pattern progression

To further elucidate the molecular heterogeneity of four histologic patterns, we obtained scRNA-seq data of six LUAD samples (one lepidic,one papillary, two acinar and two solid) from a prior study [12]. After quality control, 35,107 cells remained for subsequent analysis. We normalized and standardized the data and selected the top 2,000 highly variable genes. Utilizing the Uniform Manifold Approximation and Projection (UMAP) dimensionality reduction, we identified 21 cell clusters (Fig. S1), comprising 13 cell types (Fig. 2a). The mean expression levels of the marker genes of each cell type were shown in Fig. 2b. The epithelial cell marker genes EPCAM, KRT19, and KRT7 were used to annotate tumor cells that were further verified copy number variations [16]. Accordingly, the tumor cells exhibited higher CNVs than other types of cells (Fig. 2c and Fig. S2). Comparative analysis between histolgic patterns revealed that tumor cells in the solid pattern had higher CNVs than those in the other patterns, indicating a close link between histologic pattern progression and genomic instability (Fig. 2d). By grouping all cells according to histologic patterns, we revealed the differences in cell composition and heterogeneity in the tumor immune microenvironment. In the solid stage, the infiltration level of immune cells in tumor tissues increased (Fig. 2e), with macrophages and Tregs cells showing the most significant increase, indicating the emergence of immune inhibitory microenvironment during tumor progression. The differences in immune cell composition among different patients were observed (Fig. 2f). In summary, single-cell transcriptome revealed the molecular characteristics and heterogeneity of the tumor microenvironment during histologic pattern progression.

**Fig. 2:**
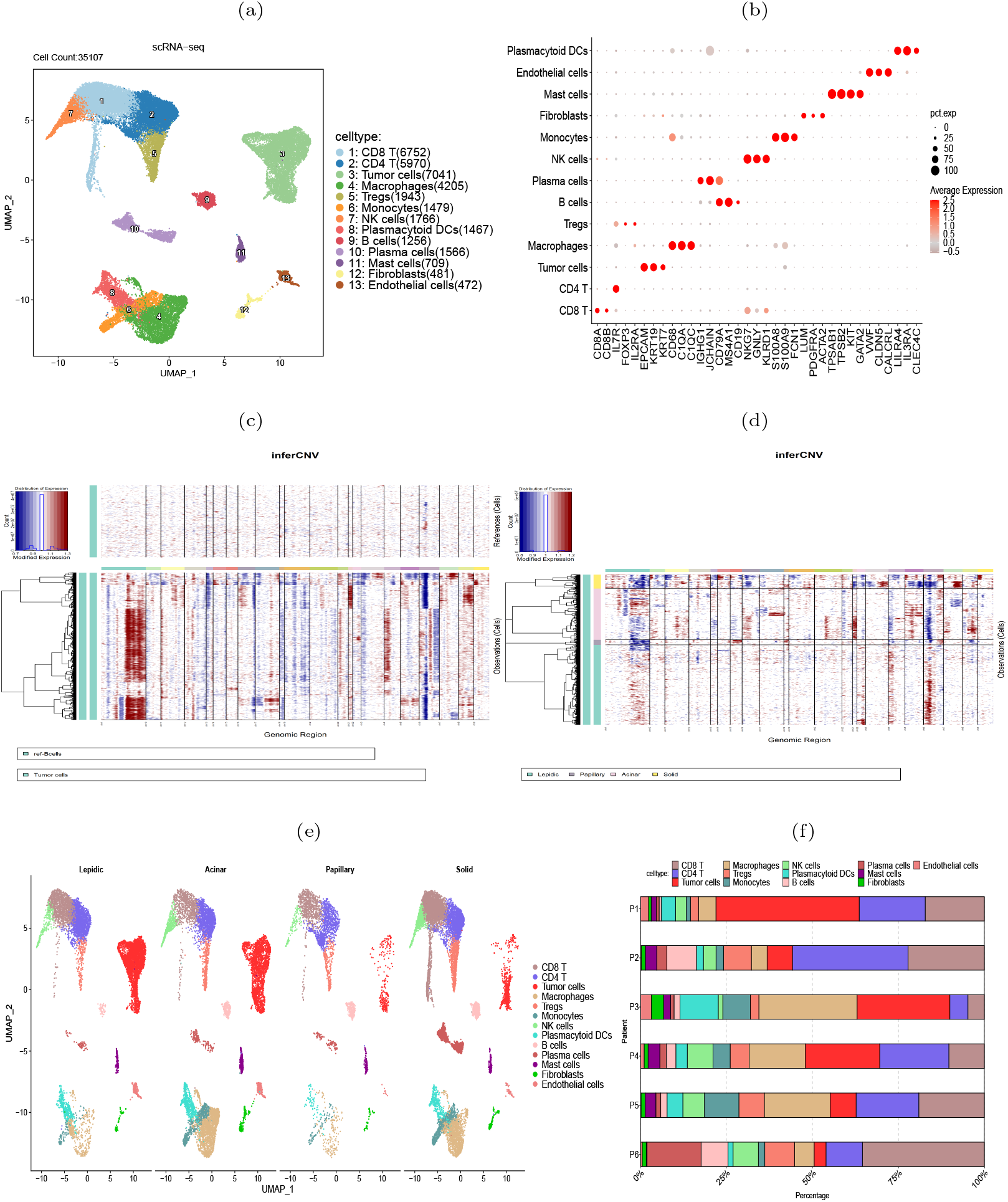
scRNA-seq data revealed heterogeneity of four histologic patterns in lung adenocarcinoma. (a) UMAP visualization of 13 major cell types. (b) Dot plots showed the average expression of known biomarker genes in specified cell clusters, with the size of the dots representing the percentage of the cells expressing the gene in each cluster. (c) Illustration of copy number alterations in tumor cells, with B cells as the reference cells. (d) Comparison of CNVs in tumor cells between four histologic patterns. (e) Cell type differences between the four histologic patterns. (f) Differences of cell compositions among three patients.

### Immunosuppressive genes increasingly expressed during histologic progression

The scRNA-seq data showed a diversity of immune inflammatory cells were recruited to the tumor microenvironment, such as tumor-associated macrophages (TAMs), lymphocytes, and mast cells (Fig. 2e). The TAMs, comprising M1 and M2 subtypes, were the predominant tumor-infiltrating immune cell population [17]. M2 macrophages mainly promote tumor growth, invasion, metastasis and immune evasion [18, 19]. We investigated the expression of HAVCR2 and LGALS9, typical immune suppressive genes, and CD163, an M2 macrophage marker gene, and found that they were co-expressed in the macrophage clusters (Fig. 3a), indicating that the TAMs infiltrating the tumor might be M2 subtype. We used gene expression profiles from the TCGA-LUAD cohort (n=598) and GSE43458 cohort (n=110) to assess the expression correlation of these gene (Fig. 3b, c). We observed that HAVCR2 gene expression had a strong correlation with CD163 (*r*=0.819 and 0.823 on two cohorts), and LGALS9 also showed weak correlation (*r*=0.292 and 0.317 on two cohorts).

**Fig. 3:**
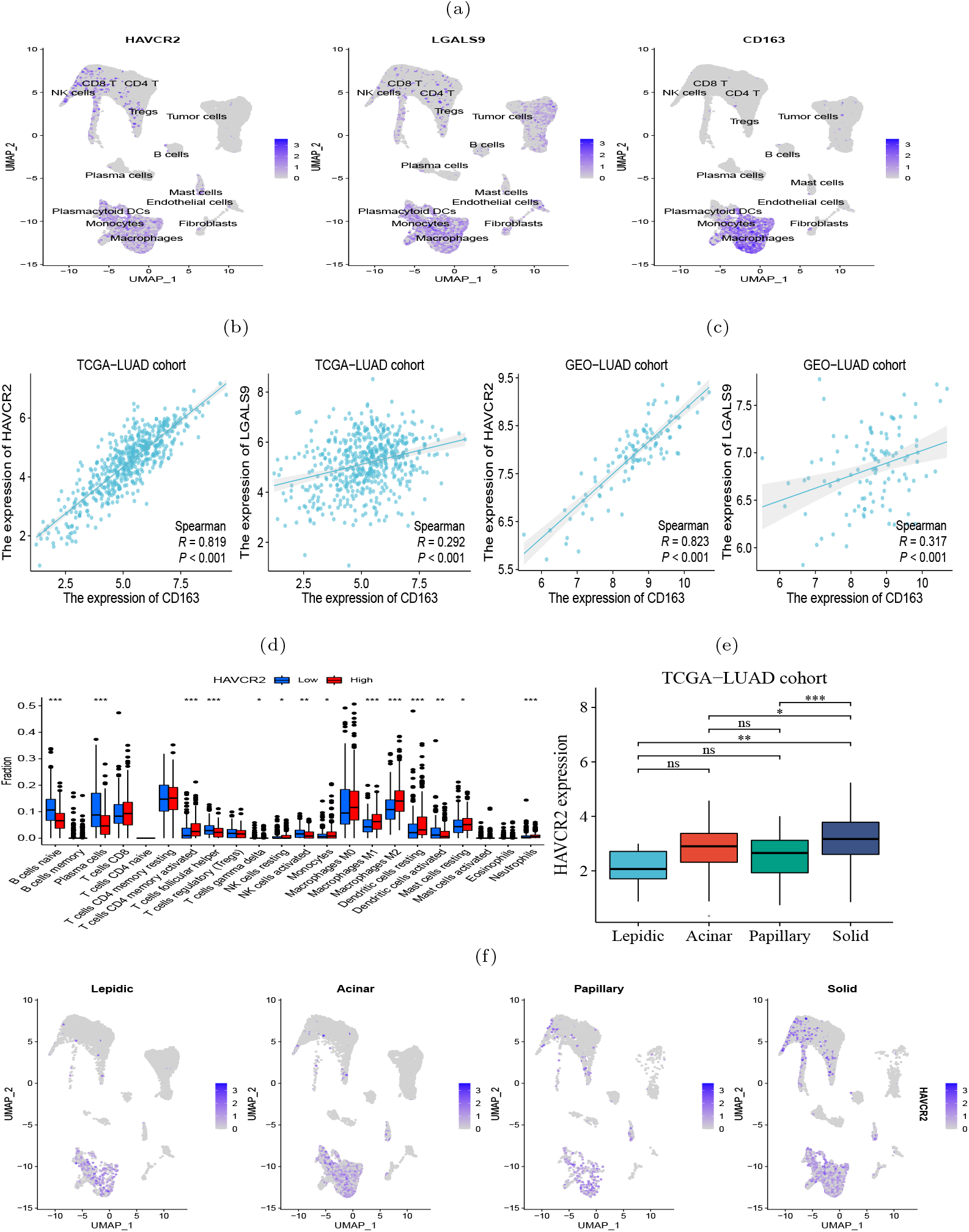
Expression of immunosuppressive biomarker genes of TAMs in four histologic patterns and their impact on immune infiltration levels. (a) Abundant expression of multiple immunosuppressive marker genes and M2 macrophage markers. (b-c) Expression of CD163 and HAVCR2 were significantly correlated in both TCGA (left) and GSE43458 (right) cohorts. (d) Difference of infiltration levels of 22 types of immune cells between two groups with high and low HAVCR2 expression. (e-f) HAVCR2 has higher expression level in solid than other histologic patterns revealed by bulk and scRNA-seq data. p<0.05 is represented by *, p<0.01 is represented by **, and p<0.001 is represented by ***.

High HAVCR2 expression in TAMs has been shown to exert immune suppressive effect [20, 21], thus we speculated that TAMs in LUAD exerted immune suppression through HAVCR2 gene. Subsequently, we stratified the LUAD patients into high and low subgroups based on the HAVCR2 expression level, and estimated the infiltration level of 22 immune cells using the CIBERSORTx tool [22]. We discovered that the M2 macrophage infiltration level in the high group was greater than that in the low group (Fig. 3d). Meanwhile, we noticed that the infiltration levels of naive B cells and plasma cells in the high group were low, which would lead to reduced ability to kill tumor cells. We further compared the expression of HAVCR2 gene in four histologic patterns. The result showed that HAVCR2 expression was higher in solid pattern than others (Fig. 3e, f), indicating that tumor microenvironment evolved to be immune suppressive during the histologic pattern progression.

### Cell-cell communication analysis revealed immunosuppressive microenvironment

During the occurrence and progression of malignant tumors, a complex cell-cell interaction network is often established to promote immunosuppressive environment and immune evasion. To study the interplay between tumor cells and other cells, CellPhoneDB [23] was used to explore the immune co-inhibitory interactions in four histologic patterns. The results revealed significant differences in the interactions among various cells between the four histologic patterns (Fig. 4a). In the lepidic pattern, tumor cells interacted substantially with macrophages and Tregs, indicating that tumor cells recruited TAMs and Tregs into the tumor at the early stage to create immunosuppressive microenvironment. With the histologic pattern progressed, the interactions between tumor cells and immune cells increased markedly. The tumor cells communicated strongly with CD4^+^ T cells, CD8^+^ T cells, Tregs, macrophages and NK cells (Fig. 4b).

**Fig. 4:**
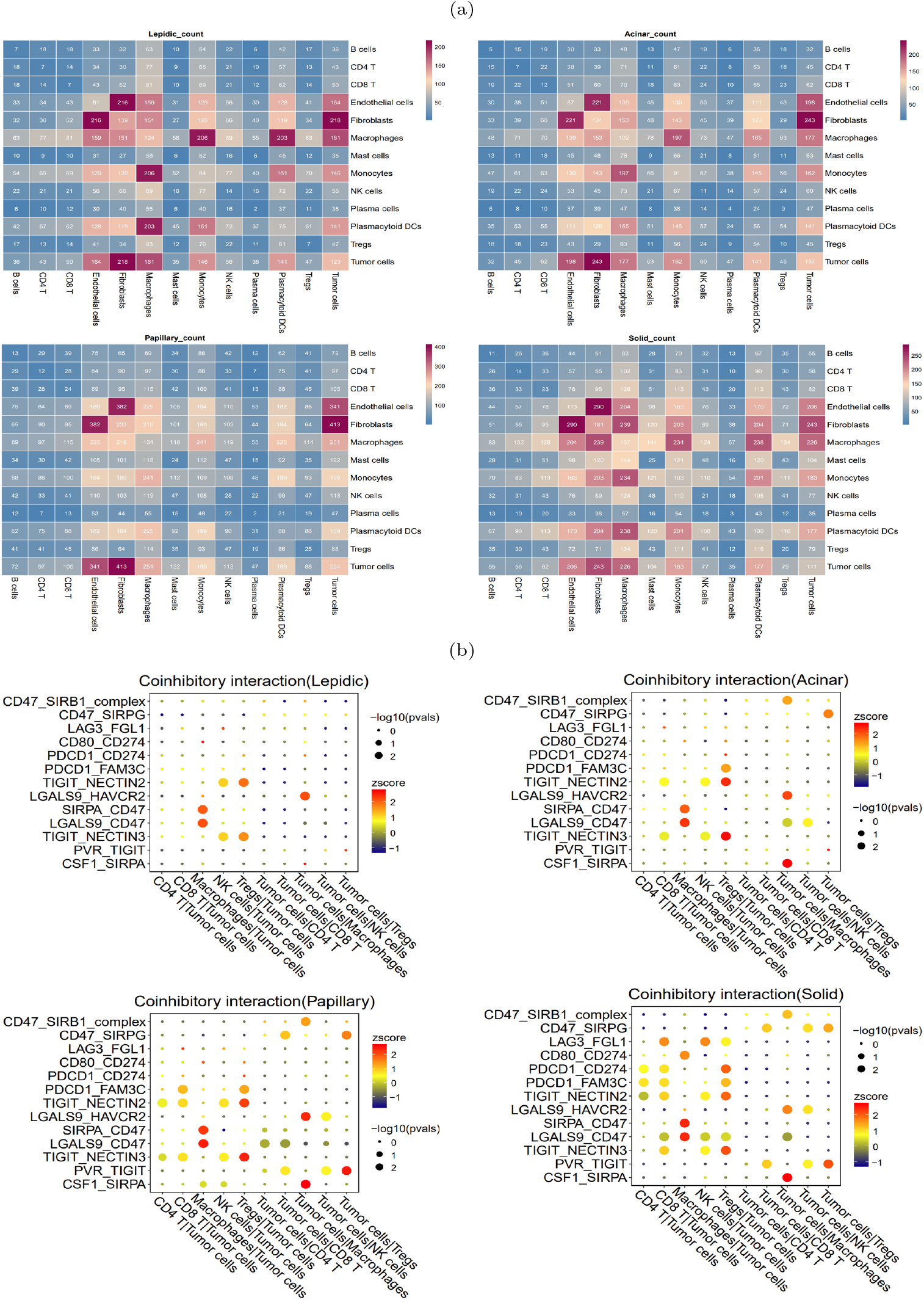
Heterogeneity of intercellular interactions in four histologic patterns. (a) Number of reciprocal pairs between cell types in four histologic patterns. (b) Co-inhibitory interactions between tumor cells and immune cells in four histologic patterns, where each row represents receptor-ligand pairs, each column represents the interacting cell types. The color and size of dots were positively correlated with the probability of intercellular interactions, with redder color representing higher expression levels (Z-scores), and larger dot size representing more significance (-log10 p-values).

In particular, we found that the NECTIN2-TIGIT immune coinhibitory interactions, mediated by the NECTIN2 ligand expressed on tumor cells and the TIGIT receptor expressed on immune cells, were prevalent during the histologic progression. In the lepidic pattern, the NECTIN2-TIGIT interactions involved tumor cells and NK and Tregs cells, and the interactions extended to encompass tumor cells and CD4^+^ T/CD8^+^ T cells with the histologic pattern progression. This suggested that the tumor cells expressed NECTIN2 to exert co-inhibitory signals and progressively inhibited an increasing number of immune cell types during tumor progression. The gene expression profiles on the TCGA-LUAD cohort corroborated that NECTIN2 was highly expressed in tumor tissues (Fig. S3). We speculated that the communicative signal between TIGIT and NECTIN2 represented a crucial mechanism for LUAD tumor cells to establish the immunosuppressive microenvironment. Indeed, several studies have confirmed that the TIGIT-NECTIN2 signal functioned to create immunosuppressive environment in other cancers. For example, single-cell sequencing and gene knockout (KO) experiments in mice have shown that the TIGIT-NECTIN2 interaction regulated the immune-suppressive environment and promoted tumor progression in hepatocellular carcinoma [24]. Xu et al. showed that the interactions occurred between the metastatic breast cancer cells and TME cells, and their spatial co-localization was verified by immunofluorescence experiments [25].

In contrast to the lepidic pattern, the solid pattern exhibited increased significance of two additional receptor-ligand interactions, PDCD1-CD274 (PD1-PDL1) and PVR-TIGIT, between tumor cells and T cells. It is well-established that the engagement of immune checkpoint receptors (PD-1) and ligand (PD-L1) is a crucial mechanism by which tumor cells evade immune surveillance [26]. TIGIT (T cell immunoglobulin and ITIM domain) is another immune inhibitory receptor protein that has been identified to be a reliably marker of T cell exhaustion [27] and a promising therapeutic target [28–30]. PVR is widely expressed in various types of solid tumors [31–34]. The immunohistochemistry results from HPA database showed that PVR is significantly enriched in tumor tissues (Fig. S4). The binding of PVR to TIGIT forms a tetramer to transmit inhibitory signals, suppressing the immune cytotoxicity of T cells and NK cells [35]. We observed that PVR-TIGIT interaction was enriched between tumor cells and Tregs in the papillary and solid patterns, implying that blockade of the PVR-TIGIT axis is a potentially effective treatment for LUAD patients with papillary and solid patterns. These finding confirmed that during tumor progression immmunosupression has been reinforced through PVR-TIGIT co-inhibitory interaction. These findings motivated us to estimate the impact of the expression levels of NECTIN2 and PVR on the tumor microenvironment and prognosis. Running the ESTIMATE tool [36] on bulk RNA-seq data from the TCGA-LUAD cohort, we calculated the level of tumor immune infiltration and found that high expression of NECTIN2 and PVR genes inhibited immune cell infiltration levels (Fig. 5a, b). We also assessed the prognostic value of these two genes based on the clinical data of TCGA-LUAD cohort and found that high expression of both genes correlated with poor patient prognosis (Fig. 5c, d). Furthermore, we analyzed the IPS immune scores of 512 patients who received anti-PD-1/CTLA-4 immunotherapy in the TCIA database [37] and found that high expression of the PVR gene significantly reduced the immunotherapy response for LUAD patients (Fig. 5e-g). Univariate and multivariate Cox regression analysis showed that PVR gene is an independent prognostic factor (Fig. S5). These results suggested that overexpression of NECTIN2 and PVR in tumor cells played an important role in the establishment of immune-suppressive environment during LUAD progression.

**Fig. 5:**
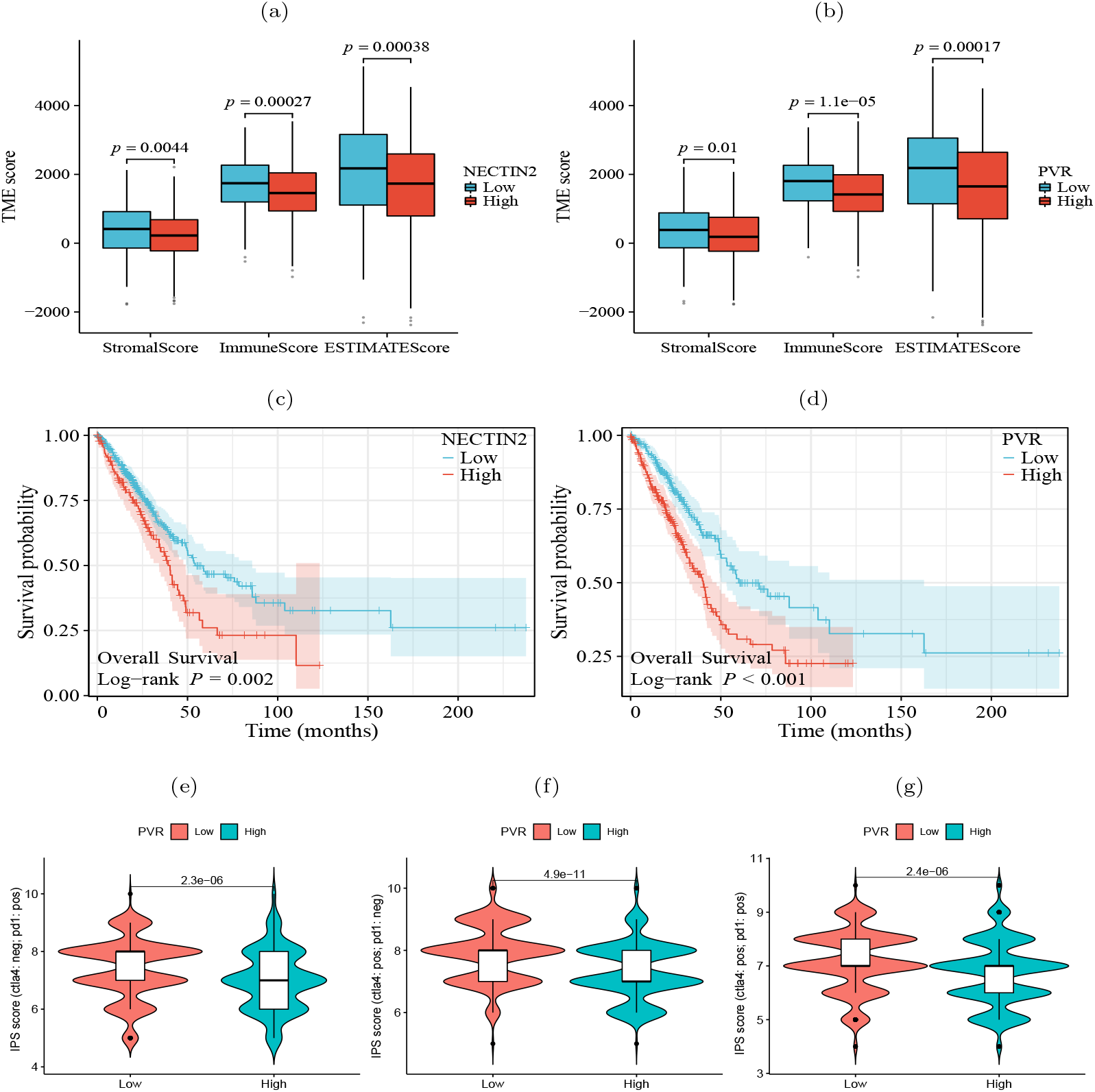
Significance of tumor ligand/receptor in immune infiltration, patient prognosis and immunotherapy. (a-b) High expression of NECTIN2 and PVR was significantly associated with poor immune infiltration. (c-d) High expression of NECTIN2 and PVR was significantly associated with poor overall survival probability. (e-g) High expression of PVR was significantly correlated with poor immunotherapy benefit. Horizontal coordinates indicates gene expression group (low and high) and vertical coordinates represents immunotherapy scores. IPS score (ctla4: neg; pd1: pos) represents the IPS score when receiving only anti-PD1 therapy, IPS score (ctla4: pos; pd1: neg) represents the IPS score when receiving only anti-CTLA4 therapy, IPS score (ctla4: pos; pd1: pos) represents the IPS score when receiving both anti-CTLA4 and anti-PD1 therapy.

### Immunofluorescence assays and spatial transcriptome validated co-localization of immune coinhibitory ligands and receptors

Our analysis above has revealed that TIGIT-NECTIN2 communication was highly active in four histologic patterns, whereas the PVR-TIGIT communication was only prominent in the papillary and solid patterns. To validate this observation, we performed immunofluorescence staining on TIGIT, NECTIN2 and PVR proteins on the pathological sections with lepidic and solid patterns. The results showed that TIGIT and NECTIN2 co-localized spatial in both types of tissues (Fig. 6a), reflecting that the tumor cells and immune cells communicated frequently through this receptor-ligand axis to create immune-suppressive environment. In contrast, PVR and TIGIT did not exhibit close proximity in lepidic tumor tissue but co-localized spatially in solid tissue (Fig. 6b). This findings suggested that tumor cells in the solid pattern enhanced immune evasion by expressing more immune co-inhibitory molecules.

**Fig. 6:**
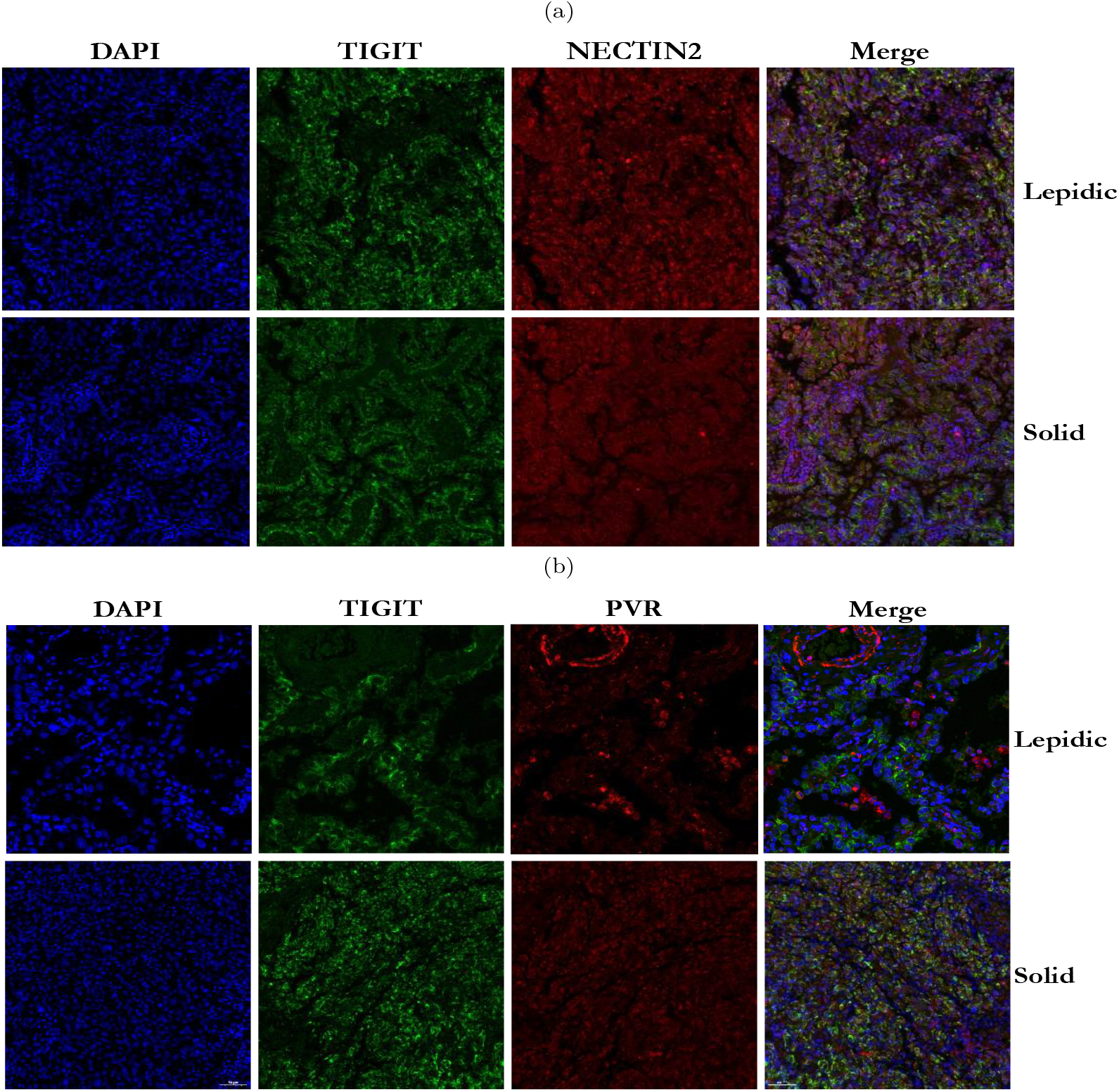
Immunofluorescence staining of TIGIT, NECTIN2 and PVR proteins in tumor tissues with two histologic patterns from two patients. (a) Immunofluorescence image of TIGIT and NECTIN2 proteins in lepidic and solid histologic patterns. (b) Immunofluorescence image of PVR and TIGIT proteins in lepidic and solid histologic patterns. (Scale bar = 50 *µm*).

We next performed spatial transcriptome sequencing (ST-seq) on two tumor samples with lepidic and solid histologic patterns. As Expected, we observed that the solid pattern had higher expression levels of these genes than the lepidic pattern and that they were predominantly localized in the tumor regions (Fig. 7a-d). Moreover, we detected spatial co-localization of TIGIT-NECTIN2 in both patterns, while PVR-TIGIT co-localization was more evident in the solid pattern (Fig. 7e). These findings further confirmed that these immune coinhibitory ligand-receptor axes may play a key role in mediating cell-cell communications leading to immunosuppressive microenvironment.

**Fig. 7:**
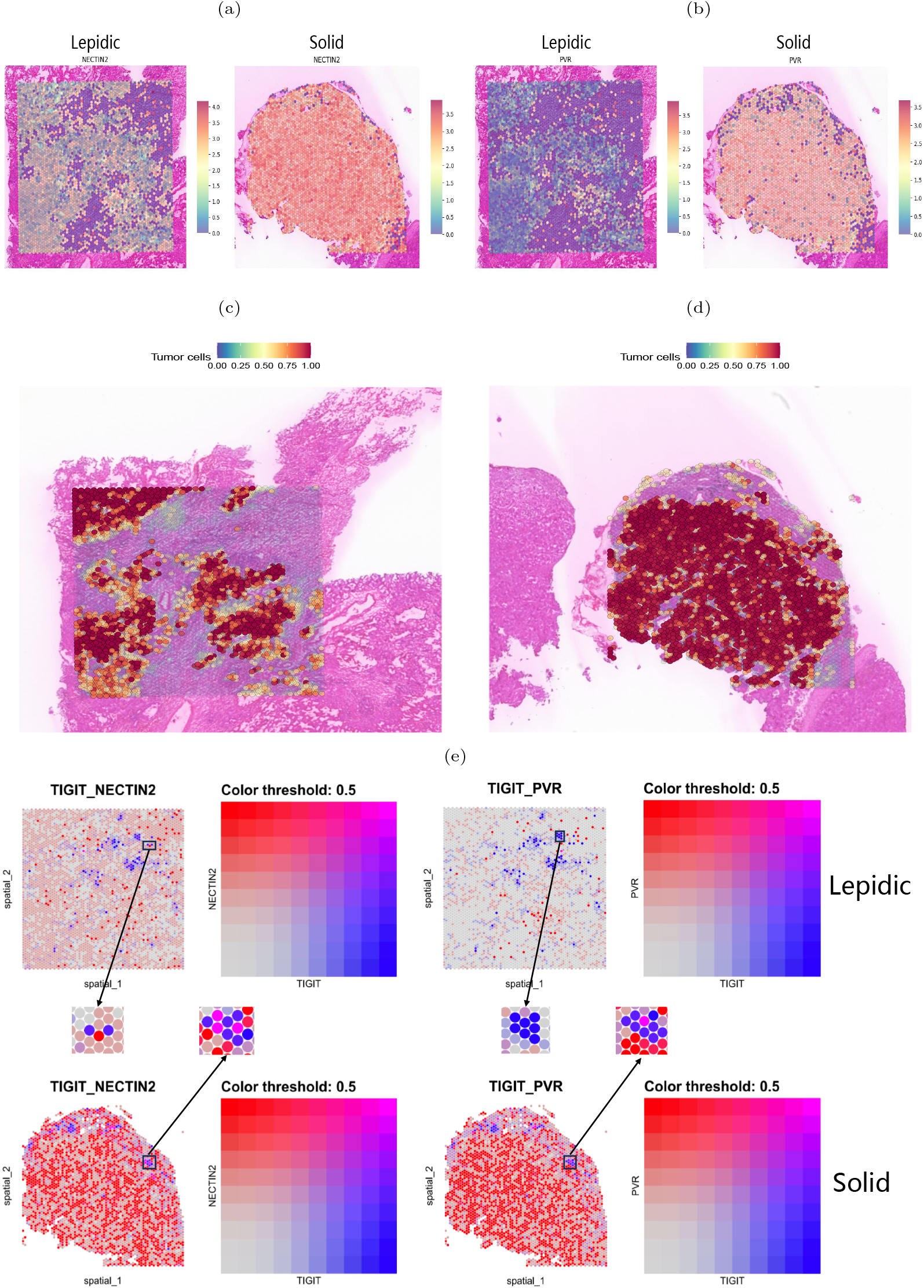
Spatial transcriptome mapping of tumor cell-associated molecule. (a-b) Expression of NECTIN2 and PVR molecules in two histologic patterns. (c-d) Tumor regions in two histologic patterns (left: lepidic; right: solid). (e) Spatial distribution of TIGIT(blue dots), NECTIN2 and PVR(red dots) in two histologic patterns. Boxes indicate adjacently expressed receptors and ligands.

### T lymphocytes transited to exhaustion during histologic progression

To verify the transition of T cells to exhaustion state during the histologic pattern progression, we clustered the CD8^+^ T cells into 13 subclusters (Fig. 8a). We examined the expression of three exhaustion marker genes LAG3, TIGIT and ENTPD1 in each cluster and found that they were highly expressed in clusters 1, 2, 4, and 13 (Fig. 8b), indicating that these T cell subclusters underwent exhaustion state transition. To compare the evolutionary trajectory association between the exhausted and non-exhausted cells, we conducted pseudotime analysis on all CD8^+^ T cells using Slingshot [38] and found that CD8^+^ T cells formed six differentiation lineages, with lineage 3, 4, 5 and 6 showing the transition from non-exhausted state to exhausted state (Fig. 8c). We also calculated the expression level changes of the three exhaustion marker genes along different differentiation lineages and found that their expression levels increased with pseudotime in the four exhaustion lineages (Fig. 8d). Moreover, we found that the three exhaustion marker genes were highly expressed in the solid pattern but almost absent in the lepidic and acinar patterns (Fig. 8e), indicating the CD8^+^ T cells transited to exhausted state in the advanced stage of LUAD.

**Fig. 8:**
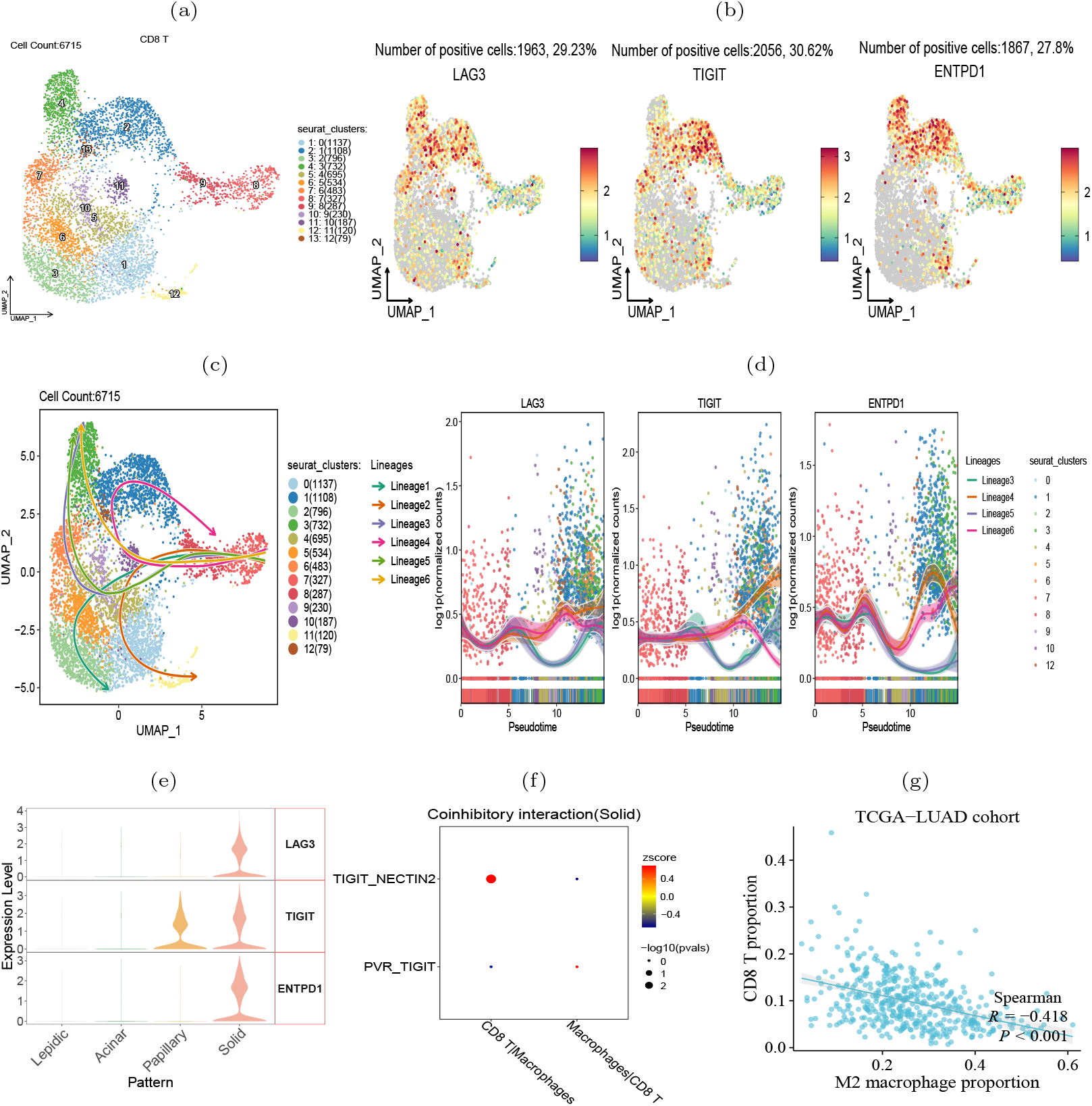
CD8^+^ T cells transition to exhaustion state during histologic progression. (a) UMAP visualization of 13 subclusters of CD8^+^ T cells. (b) Expression of exhaustion-related biomarker genes in each subcluster. (c) Illustrative plot of pseudotime lineages among subclusters, with each point representing a cell, and directed solid line the differentiation trajectory. (d) Change trends in the expression levels of exhaustion-associated marker genes over pseudotime. (e) The expression level of exhaustion-related genes in four histologic patterns. (f) TIGIT-NECTIN2 and PVR-TIGIT signals in intercellular interactions between CD8^+^ T cells and macrophages. (g) Proportion of M2 macrophages in the TCGA-LUAD cohort inversely correlated with the level of tumor infiltrating CD8^+^ T cells.

Our further intercellular communications analysis revealed a pronounced enrichment of the TIGIT-NECTIN2 interaction between the tumor-associated macrophages (TAMs) and CD8^+^ T cells within the solid tumor pattern (Fig. 8f). Previous study has demonstrated that CD8^+^ T cells interacted with TAMs to initiate the exhaustion program [39]. Accordingly, we observed that the abundance of TAMs was predictive of the scarcity of tumor-infiltrating CD8^+^ T cells, as well as reduced T cell killing activity against tumors (Fig. 8g). In summary, our analysis showed that during the progression of tumors from lepidic to solid pattern, CD8^+^ T cells gradually became exhausted and lost their ability to kill tumor cells.

## Discussion and Conclusion

In this study, we conducted comparative analysis of the tumor immune microenvironment between four histologic patterns of lung adenocarcinoma. The burden of tumor mutations gradually increases during the histologic progression from the lepidic to solid pattern. This suggested that genome transit to unstable in the later stages of tumor, leading to cellular dysfunction, disordered proliferation and poor differentiation.

The cell-cell communications landscape in the four histologic patterns indicated that as the LUAD tumor progressed, the level of immune cell infiltration increased, while the immune co-inhibitory interactions between tumor cells and immune cells synchronously intensified. In the solid pattern, there was an increase in tumor-infiltrating immune cells, especially tumor-associated macrophages and T lymphocytes. The high expression levels of exhaustion-related genes in CD8^+^ T cells reflected the immune-suppressive microenvironment in the solid pattern. Studies have reported that macrophages could hinder T cells from infiltrating into lung cancer tissue [40]. We also found that the immune-suppressive molecule HAVCR2 was not only enriched in TAMs, but also highly correlated with M2 macrophage marker gene. HAVCR2 high expression promoted the infiltration of M2 macrophages. In fact, the immune-suppressive role of HAVCR2 in hepatocellular carcinoma has been extensively studied [41, 42], but its mechanism of action in lung adenocarcinoma has not been elucidated. Our future work would explore whether HAVCR2 participate in the creation of immune-suppressive environment and anti-T cell infiltration in lung adenocarcinoma.

It was also found significant cell-cell communications between tumor cells and tumor-infiltrating immune cells via the TIGIT-NECTIN2/PVR pairs. The expression of the tumor-associated ligand/receptor genes NECTIN2 and PVR were enriched in lung adenocarcinoma and inhibited immune cell infiltration. They were also highly correlated with poorer overall survival and immunotherapy response in LUAD patients. TIGIT, a known marker gene of T cell exhaustion, is a potential therapeutic target to improve treatment efficacy for LUAD patients, via monotherapy or in combination with other drugs. For example, the phase II CITYSCOKE trial has evaluated the combination of tiragolumab (anti-TIGIT antibody) and atezolizumab (anti-PD-L1 antibody) for the treatment of non-small cell lung cancer and reported promising therapeutic effects [43]. Moreover, our immunofluorescence assay and spatial transcriptomic data confirmed the spatial co-localization of TIGIT-NECTIN2/PVR in the solid pattern.

In summary, Our integrative analysis of scRNA-seq, bulk RNA-seq and ST-seq data from four histologic patterns of lung adenocarcinoma, we have revealed the tumor heterogeneity and immune-suppressive landscape during histologic pattern progression. These findings may provide unique insights into the treatment for lung adenocarcinoma patients.

## Funding

This work was supported by National Natural Science Foundation of China (62072058, 82073339), Natural Science Foundation of Jiangsu Province (No. BK20231271).

## Contribution statement

H.L. and Q.G. conceptualized the idea. H.L. designed the experiments. H.L. and Q.G. developed the scRNA-seq and ST-seq analysis pipeline and analyzed the data. M.W. and J.L. collected clinical specimens and conducted the immunofluorescence assays. H.L. and Q.G. prepared the manuscript. J.L. revised the manuscript. H.L. and J.L. jointly supervised the research.

## Data availability

The gene expression profiles, clinical data and mutation data of TCGA-LUAD patients were obtained from https://portal.gdc.cancer.gov/. The gene expression profile of GSE43458 cohort was obtained from https://www.ncbi.nlm.nih.gov/geo/. The genomic and clinical data of MSK-IMPACT LUAD cohort were obtained from https://www.cbioportal.org/. IPS scores for lung adenocarcinoma patients were obtained from https://tcia.at/home. Spatial transcriptome data has been upload to Zenodo and is available at https://zenodo.org/records/10005621.

## Declaration of interests

The authors declare no competing interests.

## Notes

https://zenodo.org/records/10005621

### Competing Interest Statement

The authors have declared no competing interest.

